# Enabling Packaging Integration of Paper-based Electrical Gas Sensors for the Monitoring of Spoilage in Fresh Spinach

**DOI:** 10.1101/2024.07.22.604629

**Authors:** Atharv Naik, Hong Seok Lee, Jack Herrington, Giandrin Barandun, Genevieve Flock, Firat Güder, Laura Gonzalez-Macia

## Abstract

Gas sensors present an alternative to traditional off-package food quality assessment, due to their high sensitivity and fast response without the need of sample pre-treatment. The safe integration of gas sensors into packaging without compromising sensitivity, response rate, and stability, however, remains a challenge. Such packaging integration of spoilage sensors is crucial towards preventing food waste and transitioning towards more sustainable supply chains. Here, we demonstrate a wide-ranging solution to enable the use of gas sensors for the continuous monitoring of food spoilage, building upon our previous work on Paper-based Electrical Gas Sensors (PEGS). By comparing various materials commonly used in the food industry, we analyze the optimal membrane to encapsulate PEGS for packaging integration. Focusing on spinach as a high-value crop, we assess the feasibility of PEGS to monitor the gases released during its spoilage at low and room temperatures. Finally, we integrate the sensors with wireless communication and batteryless electronics, creating a user-friendly system to evaluate the spoilage of spinach, operated by a smartphone via Near Field Communication (NFC). The work reported here provides an alternative approach that surpasses traditional on-site and in-line monitoring, ensuring comprehensive monitoring of food shelf life.

## 1. Introduction

Food spoilage and waste pose formidable challenges to achieving a sustainable and efficient food supply chain. The volume of 1.3 billion tons of wasted food each year, with nearly half occurring during processing, distribution, and consumption, demands urgent attention.^1,2^

In response, there is a growing interest in developing packaging systems that go beyond containment and actively monitor the quality, storage conditions, and shelf life of food products. Monitoring gas markers of food spoilage offers a clear advantage for packaging monitoring as it does not require sample pre-treatment. For example, ammonia (NH_3_) and hydrogen disulfide (H_2_S) are marker gases for the spoilage of high-protein foods (e.g., eggs, dairy and meat) and rotting vegetables (e.g., corn and spinach).^3–8^ The integration of colorimetric, electrical, or electrochemical sensors within food packaging holds promise for achieving continuous monitoring. Current approaches in packaging integration of sensors mainly involve indicator films or on-package sensors, which provide a colorimetric response to changes in gas concentration, pH, or accumulated time-temperature history.^3,9–11^ These provide qualitative detection and visualization of freshness status. While such colorimetric sensors offer simplicity and cost-effectiveness, electrical and electrochemical sensors offer enhanced sensitivity and real-time response capabilities, enabling precise and dynamic assessment of food quality throughout the supply chain.^6,12–15^

Furthermore, integrating electrical sensors with wireless technology opens new avenues in continuous monitoring.^15–17^ With smart devices, wireless platforms comprising electrical sensors can transmit real-time data, enabling informed decisions regarding food quality control and waste reduction. Continuous monitoring systems integrating sensor technology and wireless communication facilitate traceability and enable rapid interventions to preserve food quality. Such monitoring systems consist of the following units: 1) Sensing, where molecular interaction between the target and recognition element is converted to a sensor output, 2) Decision-making, which converts raw data from the sensing unit into a human-readable format, 3) Power, which provides the supply voltage to the system by using a battery or an energy harvesting mechanism.^18–20^

To enable the widespread adoption of sensor-based monitoring systems, attention must be directed towards optimizing their integration into packaging. Encapsulation membranes enable the incorporation of sensing systems into common packaging materials without compromising food safety.^21–23^ These membranes play an important role in preserving sensor integrity while allowing efficient gas and vapor permeability, crucial for accurate detection of spoilage markers, and must achieve a balance between selectivity, stability, and physical protection.

In this work, we progress our previous research on low-cost and continuous food-spoilage monitoring based on Paper-Based Electric Gas Sensors (PEGS) to the stage of packaging integration.^24^ PEGS are near-zero-cost electrical gas sensors using cellulose paper as sensing material for the quantitative detection of water-soluble gases based on the hygroscopic nature of cellulose fibers and changes in conductivity due to dissolved aqueous species. Here we improve the long-term usage and stability of the sensors using protective membranes, paving the way for their widespread application in identifying spoilage in packed food, with a particular focus on bagged spinach. Spinach is a high-value crop, but it is prone to spoilage, with over 35% of spinach production wasted during the household consumption phase.^2^ We evaluate the performance of the sensors to monitor spinach spoilage by correlating the outputs with the microbial count of the samples. The operation of the sensors at cold (2-8 °C) and warm temperatures (24-26 °C) is also assessed to demonstrate the monitoring capacity of the system at real storage and transport conditions. The capability of sensors to distinguish between spoiled and fresh samples is demonstrated by measuring spinach samples before and after being stored in the fridge and at room temperature for several days. This system offers potential to address challenges arising from disruptions in cold chain transport or exposure to adverse environmental conditions, thus ensuring the preservation of food freshness and reducing waste. Finally, the sensor system integrated with Near Field Communication-enabled technology operated by a smartphone is demonstrated as a proof-of-concept for the wireless, batteryless detection of spoiled samples.

## 2. Results and Discussion

### 2.1 Encapsulation of PEGS

Our sensing system was based on PEGS, which has already demonstrated high sensitivity for monitoring water-soluble gases.^24^ The sensors consisted of two interdigitated carbon electrodes printed on filter paper with no other additives. A sinusoidal wave voltage was applied to the sensors at low frequency, operated by the microcontroller, and a transimpedance amplifier with a variable gain resistor was used to amplify and read the output signal (**Figure 1a**). Water soluble gases partly dissolve in water according to Henry’s law, then dissociate into ions and change the ionic strength of the solution, thereby modifying the impedance of paper.^25^

**Figure 1.**
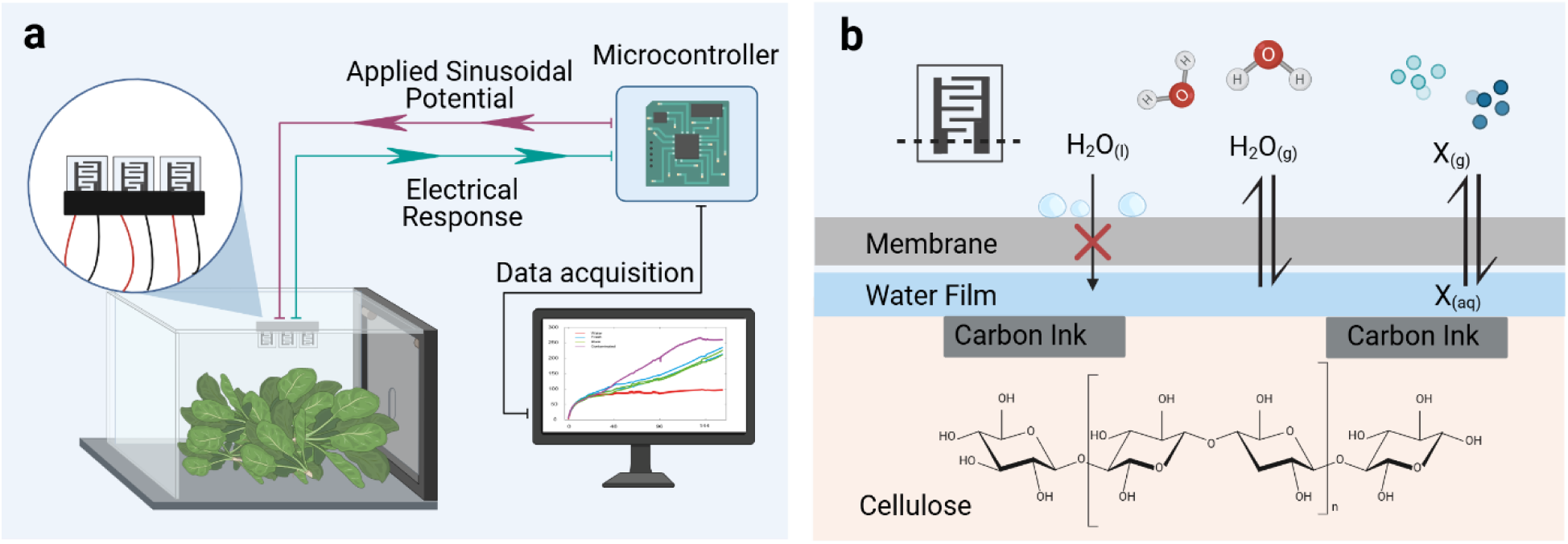
**a** Scheme of the overall system to monitor the spoilage of spinach. Encapsulated PEGS are attached to the top of the food containers, and changes in total conductance proportional to the amount of water-soluble gas released during food spoilage are monitored. **b** Schematic representation of the surface of PEGS encapsulated with thin membranes that enable the permeation of molecules in a gas state but not a liquid. The condensation of the water vapor on PEGS surface creates a thin layer of liquid covering the cellulose fibers and enabling water-soluble gases (X_(g)_) to be dissolved (X_(aq)_). Created with BioRender.com.

The encapsulation of the sensors (**Figure 1b**) represents a dual step forward towards the application of our devices in food packaging: (i) provides physical separation between the samples and the sensor, avoiding possible contamination of the electrode surfaces by food items; (ii) prevents the sensors from being wet by water condensation inside food packages, which would affect their intrinsic impedance properties.^24^ Still, PEGS need a certain relative humidity (RH >60%) to operate at its optimum, which conditions the type of material used for the encapsulation. We have tested 6 commercial membranes based on biocompatible materials such as polytetrafluoroethylene (PTFE), polyurethane (PU), polyester (PET) and cellulose. These membranes permit the permeation of gases or vapors but not liquid samples.^22,26,27^ Material specifications for each membrane tested are shown in **Table S1**. PEGS were encapsulated with the membranes using double-sided tape to seal the edges and avoid water leakages inside the sensor reservoirs (section 4.2 Encapsulation of sensors and **Figures 2a**). To evaluate the waterproofing properties of the encapsulation membranes while maintaining high RH in the sensors, encapsulated PEGS were dipped into water, and changes in conductance were recorded over time (**Figures 2b** and **S1a**). Optimal encapsulation membranes would enable changes in PEGS conductance similar to those observed by non-encapsulated sensors in a closed chamber saturated with water vapor.

**Figure 2.**
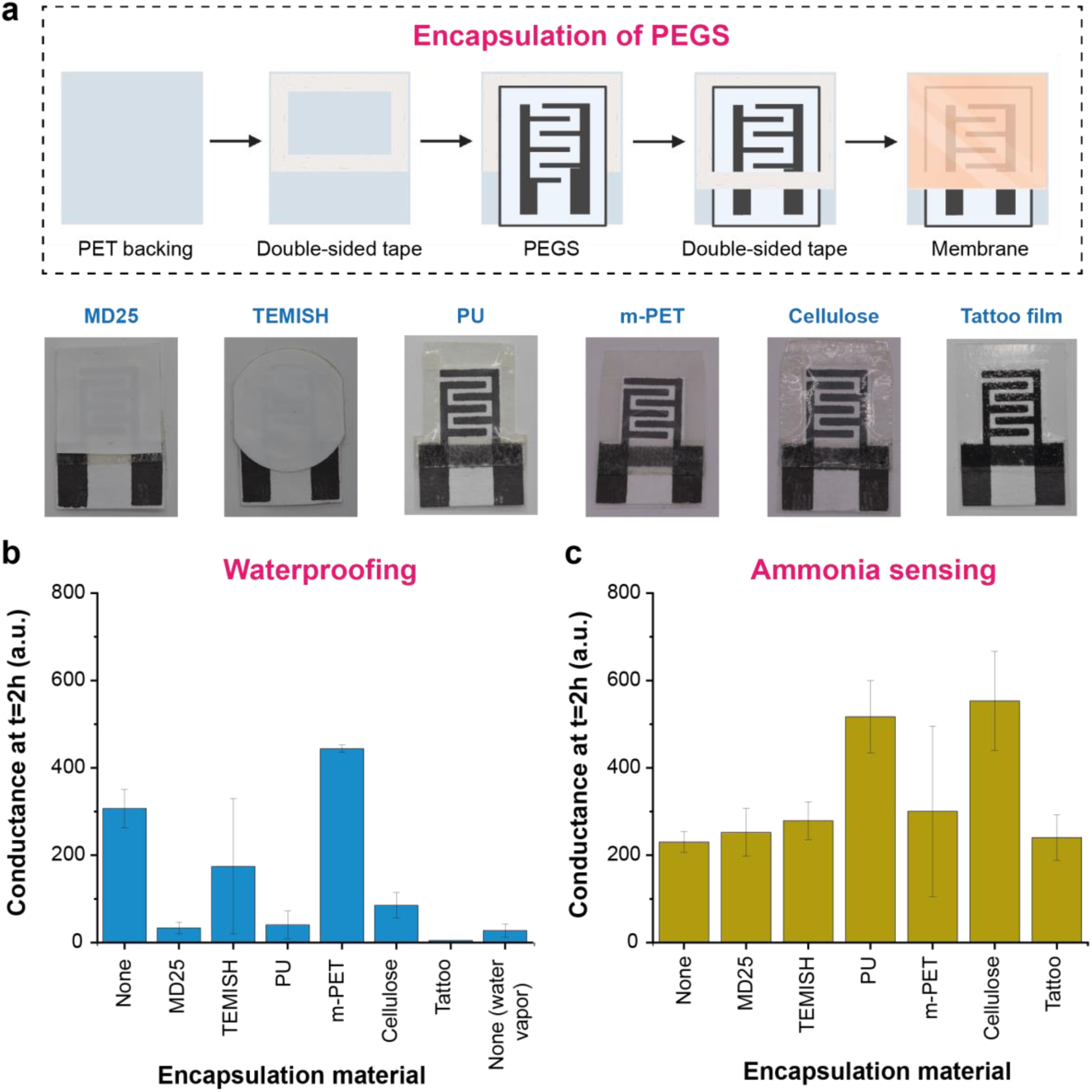
**a** Procedure followed to encapsulate PEGS with gas permeable membranes. Below: pictures of encapsulated PEGS **b** Changes in conductance of encapsulated PEGS after 2 h dipped into water and comparison with the response of non-encapsulated sensors to water and water vapor. **c** Changes in conductance of encapsulated PEGS when exposed to 1 mM NH_4_OH solution and comparison with the response of non-encapsulated sensors. PEGS are not dipped into the ammonia solution but placed above to study the determination of the gas formed in the headspace.

Only PEGS encapsulated with microperforated PET (mPET) exhibited changes in conductance higher than non-encapsulated PEGS. This could be explained by the low adhesivity of the membrane to the double-sided tape, enabling water to enter and stay in the reservoir in direct contact with the paper sensors. TEMISH membranes also showed high conductance and variability in the measurements, probably due to a decrease in the bonding properties of the adhesive layer by direct contact with water. Tattoo film membrane showed a very low response after two hours, probably due to the low permeability of water vapor. This type of polyurethane-based membranes are designed to enable optimal moisture levels of the skin while preventing water building-up under the film to reduce infection. Increasing effective permeability (Peff), solubility (Seff), and diffusion (Deff) coefficients of polyurethane films have been, however, reported with increasing RH gradient due to water vapor and polymer interactions.^28,29^ The low initial water vapor permeability might represent an issue for short response time measurements requiring rapid transduction of the response. For food monitoring, however, it is not critical as the release of spoilage gases is a relatively slow process, i.e., occurring within hours or days instead of minutes or seconds.^30,31^

We then compared the capacity of encapsulated PEGS to detect ammonia. The paper-based sensors have already shown an intrinsic selectivity towards ammonia gas in comparison to other gases tested, such as trimethylamine (TMA), hydrogen sulfide (H_2_S) or carbon dioxide (CO_2_).^24^ Encapsulated and non-encapsulated PEGS were placed in the lids of vials containing 1 mM NH_4_OH (see section 4.3 Characterization of encapsulated sensors and **Figure S1b**). Changes in the conductance of the sensors over time were due to the dissolution of ammonia gas from the headspace onto the paper-based sensors (**Figure 2c**). PEGS encapsulated with PTFE-based membranes (MD25 and TEMISH) and tattoo film showed changes in conductance similar to those registered by the non-encapsulated sensors. The changes in conductance were, however, higher when the rest of the materials were used to encapsulate the sensors, showing higher variability too. Cellulose and mPET membranes showed high water condensation inside the paper-based sensor reservoir, which could explain the high changes in conductance.

Overall, MD25 and tattoo film were optimal for the encapsulation of PEGS. They did not allow the permeation of liquid water, and their response when exposed to 1 mM NH_4_OH solution was similar to that from the non-encapsulated sensor. We then studied further applications of the sensor system in the monitoring of food spoilage.

### 2. 2 Monitoring spinach spoilage using non-encapsulated PEGS

The spoilage of fresh spinach was first monitored using raw PEGS without any encapsulation layer. The scheme of the experimental setup is shown in Figure 1.

Changes in the conductance of the sensors associated with the spoilage of spinach were monitored over a week at room temperature (25 °C). Microbial plates were run in parallel to evaluate the correlation between PEGS response and microbial content on the samples. PEGS response increased continuously during the first three days, which was attributed to an increase in the concentration of water-soluble gases released by the spinach samples during the spoilage mechanism (**Figure 3a**). The concentration of ammonia and Volatile Organic Compounds (VOCs) such as ethanol, methanol, and organosulphur compounds (like dimethyl sulphide and methanethiol) has already been shown to increase significantly in the headspace of packaged baby spinach leaves when stored for 5 days at 21 °C.^32^ PEGS has already shown high sensitivity for the detection of ammonia and, to a less extent, to other gases such as CO_2_ and trimethylamine (TMA).^24^ The difference in sensitivity of PEGS towards several water-soluble gases was explained by the different levels of dissociation, solubility, and ion mobility of the gases in water. The overall conductance of the sensors in the food containers increased from 250 a.u. (t = 24 h) to 650 a.u. (t = 72 h) whereas the sensors in the water containers (blank) were stable at approx. 85 a.u. for the entire analysis. After 72h, the conductance of the paper-based electrodes in the presence of spinach reached a plateau, showing an overall conductance almost 8-fold higher than the blank sensors. This behavior was in line with the increase of microbial colonies in the food samples (**Figure 3b**). Aerobic Colony Count (ACC) is a food quality indicator that provides information about the remaining shelf-life of a product or possible issues during handling and storage.^33–36^ Total ACC increased from 10^6^ CFU g^-1^ (t = 0 h) to over 10^8^ CFU g^-1^ (t = 48 h), the threshold for ACCs at which ready-to-eat food products such as salad vegetables are considered spoiled, reaching a plateau.^37,38^

**Figure 3.**
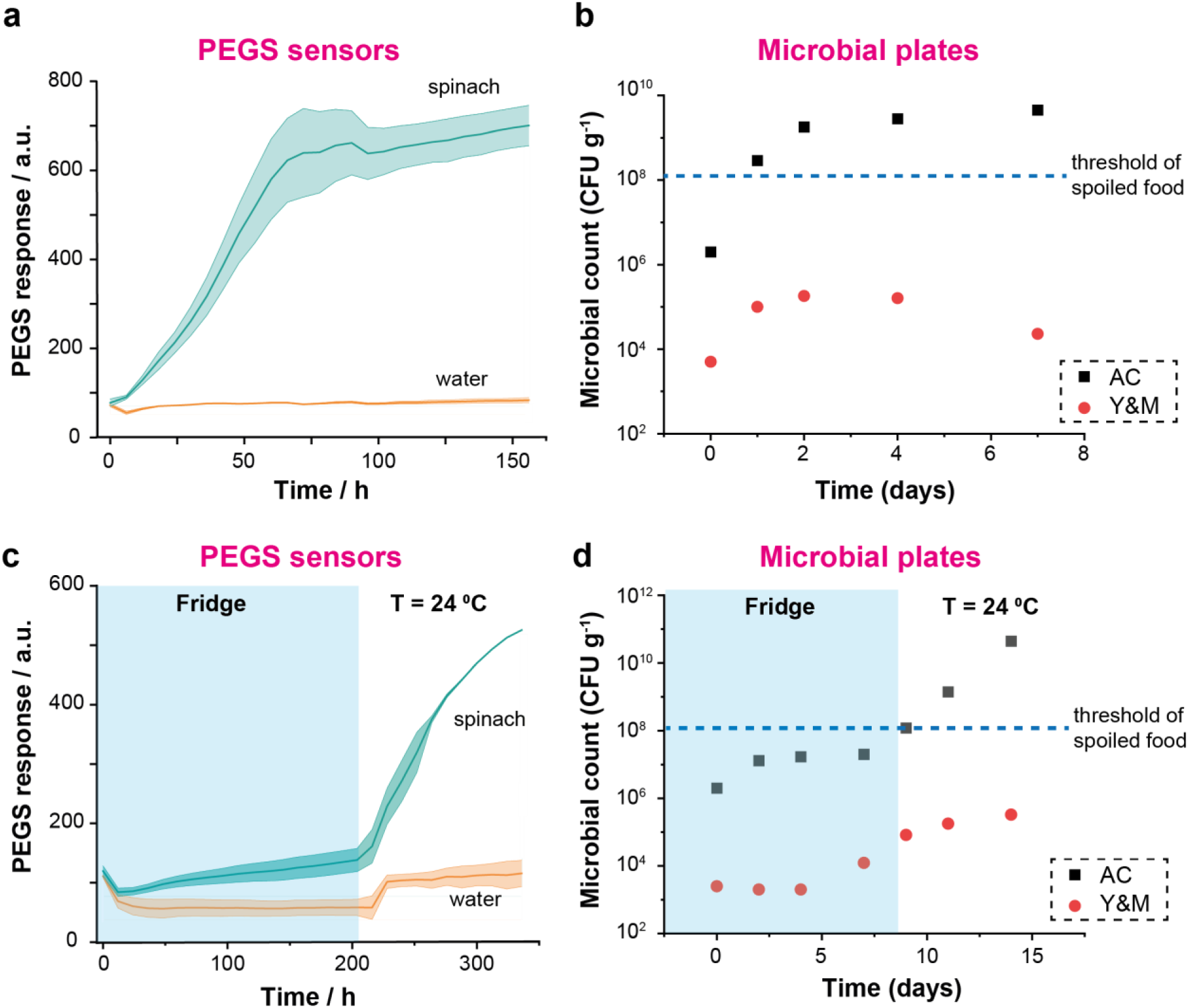
**a** Response of PEGS to spinach spoilage monitored over 6 days at room temperature (T = 25 °C, n = 3 for water and n = 4 for spinach). **b** Microbial count (ν Aerobic Count (AC) and λ Yeast and Mould (Y&M)) for the samples in a. **c** Response of PEGS to spinach spoilage monitored for 9 days in the fridge (T = 2 – 8 °C) followed by 5 days at room temperature (T = 25 °C, n = 3 (water) and n = 4 (spinach)). **d** Microbial count (ν AC and λ Y&M) for the samples in c.

Storage at low temperatures normally reduces the rate of bacterial growth and extends food shelf-life.^30,31,39^ We monitored the spoilage of fresh spinach at low temperatures using raw PEGS (without encapsulation). The correlation between the increase of PEGS signals and the quantity of bacteria in the spinach samples was also observed at low temperatures. **Figures 3c** and **3d** show the change in conductance of PEGS and the bacterial activity, respectively, when spinach and blank samples were kept in the fridge (T = 2 – 8 °C). PEGS in spinach containers showed a reduced response at low temperatures compared to room temperature, with only 2-fold higher response than blank sensors after 216 h compared to the 8-fold difference observed at T = 25 °C. This was in agreement with the data obtained from the microbial plates. The concentration of aerobic bacteria in spinach samples was constant for over one week when refrigerated, only increasing over the quality threshold after the food samples were placed at room T for a further two days.

Overall, PEGS showed a distinct change in conductance during the spoilage of spinach, in good correlation with the microbial plates, demonstrating their potential for monitoring food freshness by integration into packaging systems.

### 2.3 Monitoring spinach samples with encapsulated PEGS

We then studied the effect of PEGS encapsulation on monitoring the spoilage of spinach. **Figure 4a** compares the impact of encapsulation membranes on the sensitivity of PEGS to detect the spoilage of spinach stored in the food boxes after 7 days at T = 25 °C. The changes in the conductance of encapsulated PEGS when exposed to only water are also shown as controls. Based on these results and their performance during the waterproofing and ammonia sensing tests (section 2.1), we selected the most suitable membrane to encapsulate PEGS towards their integration into food packaging. The criteria we followed were: *(i)* high waterproofing, *(ii)* sensing properties towards ammonia and spinach (similar or better than non-encapsulated PEGS), and *(iii)* cost.

**Figure 4.**
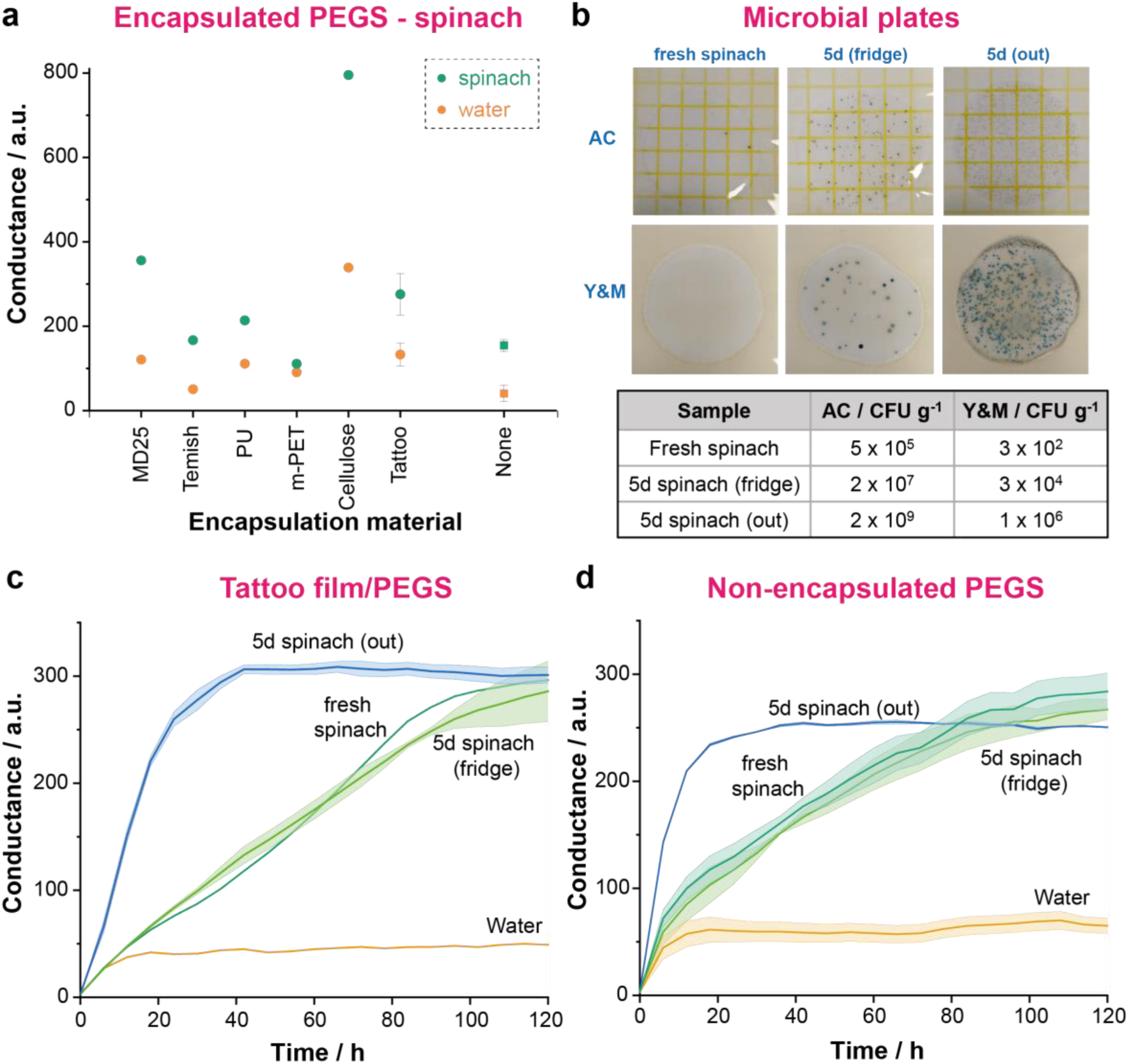
**a** Comparison of changes in conductance of PEGS encapsulated with different membranes and non-encapsulated PEGS (none) exposed to food boxes containing spinach (sample) or water (control) after 7 days (none: n = 3 water, n = 6 spinach; tattoo: n = 2 water, n = 4 spinach; other materials: n = 1). **b** Top: Microbial plates at 1:10^5^ dilutions (AC) and 1:10^3^ dilutions (Y&M) of fresh spinach, 5-day-old spinach stored in the fridge and 5-day-old spinach stored out of the fridge at T = 25 °C (all spinach bags were opened within 2 h of measuring the microbial content). Bottom: Table with microbial content in same samples, measured on the first day of the experiment (D0). Dynamic curve of PEGS conductance within food boxes containing water, fresh spinach, 5-day-old spinach stored in the fridge or 5-day-old spinach stored when PEGS are: **c** encapsulated with tattoo film (n = 1, water and fresh spinach; n = 2, 5d spinach fridge and out) and **d** non-encapsulated (n = 2).

m-PET did not show any difference between water and spinach sensing. This membrane had already shown high variability in ammonia monitoring (Figure 2b) and lack of waterproofing with the current encapsulation approach (Figure 2a), making it unfeasible for this application. mPET behavior might be explained by the loss of adhesivity of the membrane to the double-sided tape after long exposure to a high-humidity environment. This accumulated water in the reservoir, in direct contact with the paper sensors, which might potentially change the sensing mechanism. Additionally, PET has already been reported to be degraded by some of the gases generated during spoilage gases (e.g., the combination of ammonia and CO_2_), which might affect the reproducibility and sensing capabilities of PEGS.^40^

Cellulose showed high responses to both spinach spoilage and control water, translating into an overall higher sensitivity than non-encapsulated PEGS. This membrane also showed high water absorption, which was expected due to its cellulosic nature, leading to the accumulation of water in the PEGS reservoir. This might explain the high responses to the gases and the high variability observed in ammonia sensing. PEGS are paper-based, and their continuous contact with water might degrade the sensors over time. For this reason and despite its potential, cellulose membrane was not selected for further long-term studies.

MD25 showed high potential as an encapsulation membrane with good waterproofing and sensing capabilities for ammonia and spinach spoilage. Its cost, however, makes it unsuitable for the encapsulation of low-cost PEGS as it would increase its total price by at least 20-fold.^24,41^

By comparing PEGS response to the spoilage of spinach with respect to water, PU, TEMISH and tattoo film showed responses similar to those shown by the non-encapsulated sensors. PU, however, showed some disruptions in the encapsulation sealing after prolonged exposure to moist environments. This was not noticeable at short times, for example, after dipping for 2 h in water (Figure 2a), but it was observed in the sensors exposed to the spoilage of spinach for 7 days. Like MD25, TEMISH is PTFE-based, and its high cost makes it unsuitable for this application. For these reasons, PU and TEMISH were discarded and only tattoo film was considered for further experiments towards integrating PEGS into food packaging.

We then tested the capability of encapsulated PEGS (tattoo film) to differentiate spoiled and fresh spinach. For this investigation, we used two bags of spinach from the same batch; one was stored for 5 days in the fridge (*5d spinach (fridge)*), and the other one was kept outside the fridge at 25 °C (*5d spinach (out)*). Fresh spinach purchased on the starting day of the test and water were both used as controls. Food boxes containing 20 g of each sample or 100 g of water were monitored over 5 days using encapsulated PEGS (tattoo film) and non-encapsulated. The microbial content (AC and Y&M) of the spinach samples at the start of the test was also recorded (**Figure 4b**). Spinach samples kept outside the fridge for 5 days showed a high level of AC and Y&M content, over the threshold considered for food safety. PEGS also showed a remarkable increase in the response when exposed to *5d spinach (out)*, which agreed with the results from the microbial plates (**Figures 4c and 4d**). PEGS response then reached a plateau after 30-40h. The response of PEGS for both *5d spinach (fridge)* and fresh samples was similar. They both showed a mild but continuous increase of PEGS signal over time caused by the spoilage of the samples. The 5-day-old sample showed higher initial microbial content but still within the quality threshold, both samples being indistinguishable from each other by the naked eye. Samples stored at low temperature had already shown a slow spoilage rate and, therefore, PEGS response (Figures 3c and 3d). This agrees with PEGS responses for *5d spinach (fridge)* and fresh samples. Both signals reached a plateau after 100-120h of monitoring, at the same level as *5d spinach (out)*. This is consistent with the standard pattern of bacterial growth in closed systems, which determines the VOCs released during food spoilage.^42,43^ The encapsulation of PEGS with tattoo film did not seem to hinder their capacity to monitor the spoilage of spinach. Encapsulated PEGS showed a slight delay in the response in the first hours of the test since the presence of the membrane probably slows down reaching the RH value required for their optimum operation. This delay, however, did not interfere with the ability of the encapsulated sensors to provide an accurate pattern of spoilage.

### 2.4 NFC Integration

We then demonstrated the capacity of encapsulated PEGS to monitor food spoilage via integration into food packaging. We designed a disposable NFC-powered device for batteryless and wireless monitoring of food freshness on-site. **Figures 5a** and **5b** show the system architecture and design of the device, consisting of a planar single-coil copper antenna with a resonant frequency of 13.56MHz. The SiliconCraft SIC4341 type 2 tag IC with a potentiostat sensor interface was used to obtain conductance readings and communicate with the smartphone simultaneously. Through a custom smartphone app, the IC was programmed to produce a 1.6 V peak-to-peak and 5 Hz square wave signal across the PEGS, and the conductance was measured over 20 seconds. The device was fabricated as a flexible PCB, with a ‘window’ to expose PEGS, and then encapsulated using polyurethane-based tattoo film (the pre-existing adhesive layer on the tattoo film also enabled mounting the device within spinach packages). By using low-cost, commercially available materials and electronic components we proved the viability for widespread adoption.

**Figure 5.**
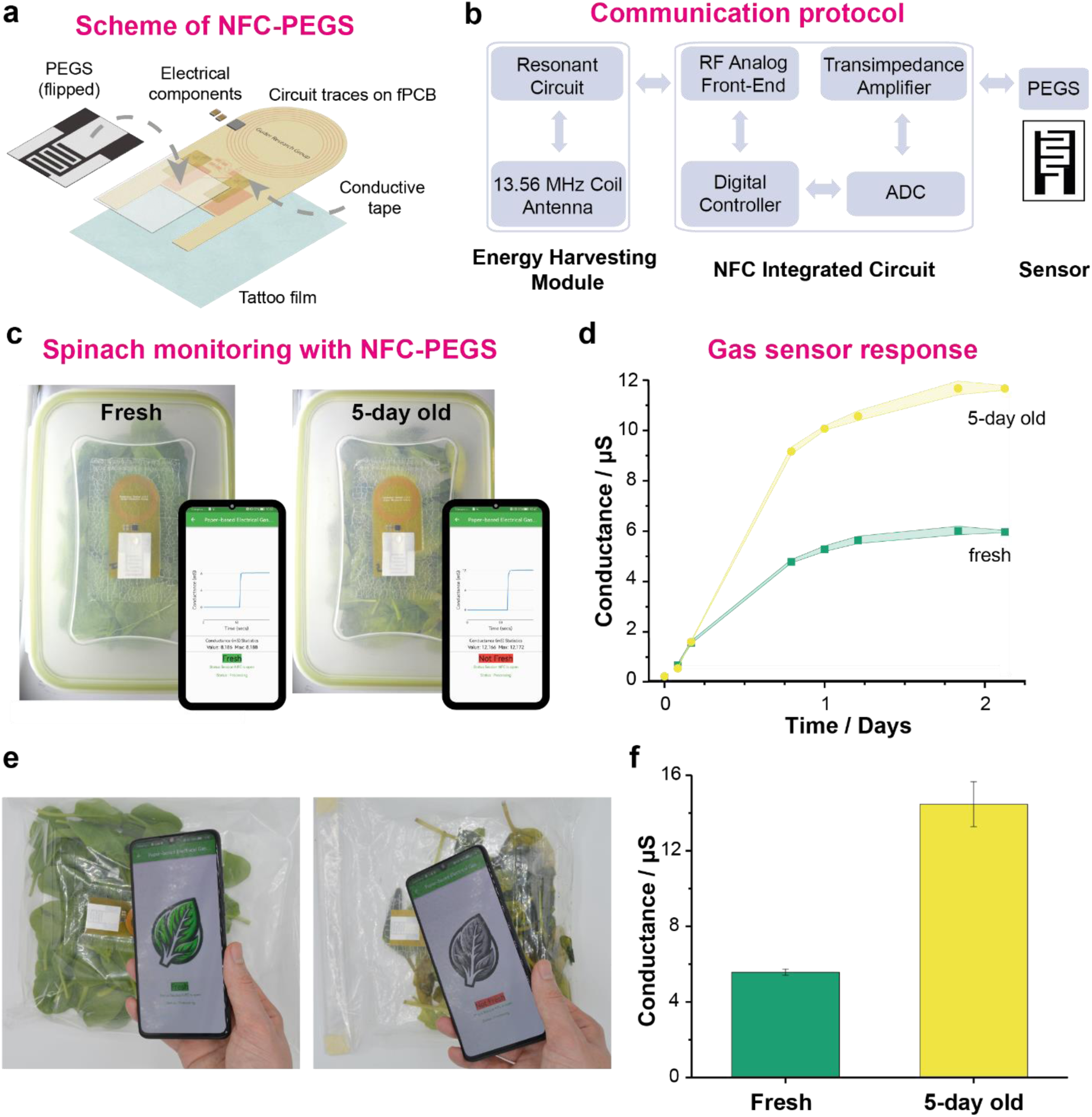
**a** Schematic illustrating the exploded view of the NFC-powered batteryless device including fPCB, PEGS and encapsulation. **b** Block diagram illustrating the key components comprising the NFC electronics of the device. **c** Pictures of the NFC-powered sensing device with encapsulated PEGS integrated into food boxes containing fresh (left) and 5-day old (right) spinach. Inset: Pictures of the app developed (research version) to run the measurements, plot the conductance, and communicate the *fresh* or *not fresh* result. **d** Conductance of PEGS over time within the food boxes containing fresh and 5-day old spinach recorded by the NFC-devices and smartphone app (n = 4). **e** Pictures of the NFC-powered sensing device with encapsulated PEGS integrated into polyethylene food bags containing fresh (left) and 5-day old (right) spinach and smartphone app (user-friendly version) showing the *fresh* or *not fresh* result of the measurement. **f** Comparison of PEGS conductance from (e) recorded after 24 h (n = 4).

Figure 5c is a demonstration involving the NFC-powered devices placed within food boxes containing *i)* fresh spinach (purchased on the day of the measurement) or *ii)* 5-day-old spinach (unopened bagged spinach stored at 25 °C for 5 days). The real-time freshness of each package was determined based on the conductance reading of the PEGS, with thresholds determined from prior experiments, and communicated to the smartphone. We used a research version of the app, which showed the real-time conductance on-screen during the measurement (20 s) after tapping on the NFC-PEGS device. The data stored in the phone were then transferred to the cloud and used to plot the conductance readings (Figure 5d). As expected, data from both samples were indistinguishable for the first few hours until the RH reached the optimum value for operation. After that, the system allowed the user to differentiate between fresh and spoiled spinach samples by a 2-fold conductance difference. The repeatability of the NFC-PEGS device and app operation was proven by recording subsequent readings (n = 4) of each sample at each time point, showing relative standard deviations (RSD) below 5% for all the measurements.

We then demonstrated the potential of the NFC-PEGS tags to monitor food freshness by integrating the device into spinach packages of varying levels of freshness (Figure 5e). 40 g of i) fresh spinach or ii) 5-day-old spinach were placed in polyethylene zip bags with the NFC-PEGS tag attached on the side. The samples were stored at 25 °C and measured over 3 days. A user-friendly version of the app was used here to communicate the level of spinach freshness by “Fresh” or “Not Fresh” messages. The research app was also used to collect the data and build the comparative graph (Figure 5f). The intra-assay repeatability, measured by subsequently recording (n = 4) each sample at each time value, was below 10% RSD.

## 3. Conclusion

We have presented here a system for monitoring spoilage in packed foods based on Paper-Based Electric Gas Sensors integrated within packaging through gas-permeable membranes. This allows the permeation of analyte gases/vapors while preserving the stability of the sensors and protecting integrated electronics through waterproofing. Among the membranes we tested (MD25, TEMISH, PU, m-PET, cellulose, and tattoo film), the polyurethane-based tattoo film showed an optimal balance between response and waterproofing, with performance comparable to non-encapsulated sensors.

This work provides a standard method for using encapsulated PEGS that does not require highly specialized equipment and can be performed in most labs. As a proof-of-concept, we demonstrate NFC-enabled wireless, batteryless detection of spoilage in spinach. Given the efficacy and low cost of the full system (US $0.35), it could be implemented within food packages to enable dynamic assessment of spoilage beyond traditional expiry dates, in a disposable manner. The current packaging integration, however, has the following three challenges:

i. In testing membranes, where a membrane did not have a pre-existing adhesive layer, double-sided tape was used to attach the membranes onto PEGS. This made it difficult to judge long-term response in case of a decrease in the adhesive bonding over time; gases may penetrate the device via areas of reduced bonding rather than through the membrane.
ii. In this work, microbial counting was used to evaluate the level of spinach spoilage detected by the encapsulated sensors as a consequence of the overall amount of gases released. For further studies exploring spoilage mechanisms in detail, instrumental analysis techniques like gas chromatography-mass spectrometry (GC-MS), selected-ion flow-tube mass spectrometry (SIFT-MS) and secondary electrospray ionization mass spectrometry (SESI-MS) may be required for gas identification and quantification. This would help researchers understand the differences in spoilage mechanisms and gas equilibrium occurring inside food boxes compared to bags, which would explain the high inter-assay variability observed when NFC-PEGS tags were tested in polyethylene zip bags.
iii. Additional modifications of the paper-based sensors (e.g., with water, acids, and hydrogels) have been previously used to accelerate the detection of spoilage by PEGS.^24,44^ Here, only unmodified dry sensors have been used, which take longer to stabilize, thereby losing the initial response. Since our platform will be integrated into packed spinach at the packaging stations, the sensors will have time to equilibrate before the spoilage process starts. For future applications, modifications like hydrogels may enable the capture of the initial response by providing faster stabilization.

In this work, we focused on applications in spinach spoilage but our system is flexible and can be easily applied to other packed foods or perishables. Further research on marker gases/VOCs released over spoilage of various foods and their permeation characteristics through encapsulation membranes can be used to optimize parameters such as membrane thickness and headspace within the encapsulated device. Despite this study’s focus on PEGS, the encapsulation scheme presented can easily be applied to other sensors. For example, encapsulation membranes can be used to create a thin layer of electrolyte near the sensors, where electrochemical reactions can be studied. With a broader scope, development of systems capable of simultaneous powering and communication with NFC tags in multiple packages (through anti-collision protocols) can enable continuous monitoring of freshness throughout the supply chain.

## 4. Methods

### 4.1 Reagents

Monobasic potassium phosphate, ammonium hydroxide (NH_4_OH) and sodium hydroxide were purchased from Sigma-Aldrich, UK.

The paper-based electrical gas sensors (PEGS) were screen printed using 90% conductive carbon sensor paste (C2030519P4, SunChemical) and 10% thinner (CDSN4059, SunChemical) by Calder Screenprint Ltd, UK. The electrodes were printed on Whatman^TM^ grade 1 chromatography paper (20 cm × 20 cm, 0.18 mm thickness). We used the same size and configuration as previously optimized in our group.^24^ POREX® Porous PTFE medical material (MD25) and 3M™ Polyurethane medical film 9832F (PU) samples were kindly provided by Parafix, UK. TEMISH® porous PTFE S-NTF8031J samples were kindly provided by Nitto, Japan. Biaxially oriented polyester film (OCLF, mPET) and cellulose-based compostable sealing film (cellulose) samples were kindly provided by Bullseye Food Packaging, UK. Polyurethane-based tattoo film (Tattoo) was purchased from Amito E-commerce Co., LTD, UK. 3M™ 9086 Translucent double-sided tape (0.19mm thick) was purchased from RS Components, UK. 3M™ Petrifilm™ Aerobic count (AC) plates and 3M™ Petrifilm™ Yeast and Mould (Y&M) count plates were purchased from Scientific Laboratory Supplies, UK.

### 4.2 Encapsulation of sensors

The procedure followed to encapsulate PEGS with the membranes under study is indicated in Figure 2a. Briefly, double-sided tape (3M™ 9086, 0.19mm thick) was used to fix PEGS to a PET substrate and seal the edges, delimiting the sensing area and avoiding water leakages inside the sensor reservoirs. After placing the paper-based sensors, an extra layer of tape was added to the front part of the sensor to ensure the sensor reservoir was fully sealed once the encapsulation membrane was brought into contact. After encapsulation, the sensor reservoir was 1.2 cm x 1.8 cm. The encapsulation membranes covered only one side of the sensors to facilitate the study.

### 4.3 Characterization of encapsulated sensors

#### Waterproofing

Encapsulated PEGS were placed in 28-mL vials containing 25 mL of deionized (DI) water, leaving the sensors dipped into water (Figure S1a). Unless otherwise stated, changes in PEGS conductance were then recorded by applying a sinusoidal voltage signal with an amplitude of 4 V and 10 Hz to the sensors and using a transimpedance amplifier with gain resistor of 50 kΩ to amplify and read the output signal (current). We measured the amplitude of this signal, which corresponded to the magnitude |Z| of the impedance of the sensor. Measurements were taken every 2.5 s.

#### Ammonia sensing

Encapsulated PEGS were placed in 28-mL vials containing 10 mL of 1 mM NH_4_OH, where the sensors were not in contact with the solution but in the headspace (Figure S1b). Changes in PEGS conductance were recorded using a sinusoidal voltage signal of 4 V and 10 Hz for at least 24 h.

### 4.4 Monitoring spinach spoilage

For sample monitoring, 20 g of fresh bagged spinach was placed in 1L food containers with PEGS attached to the lids. Additional containers with 100 g of DI water to create 100% RH were used as controls. Each container accommodated two sensors. Each experiment consisted of six plastic containers (12 sensors evaluated in total), four with spinach and two with water as controls. For experiments involving non-encapsulated PEGS, 20 µL of DI water were deposited onto PEGS immediately before placing the sensors into the food containers to enhance signal recording.^24^ For experiments with encapsulated PEGS, they were used dry from the start to mimic food packaging procedures. Unless otherwise stated, a sinusoidal voltage signal with an amplitude of 2 V and 10 Hz was applied to the sensors, and a transimpedance amplifier with gain resistor of 50 kΩ was used to amplify and read the output signal.

### 4.5 Microbial plate counting

Potassium dihydrogen phosphate stock solution was prepared by adding 34 g of monobasic potassium phosphate and 175 mL of 1 N sodium hydroxide to 825 mL of deionized water. Butterfield’s buffer (pH 7.2) was prepared by adding 1.25 mL of potassium dihydrogen phosphate stock solution to 1 L of deionized water, followed by sterilization.^45^ To measure the microbial content of spinach samples, 10 g of spinach were first blended with 90 g of Butterfield’s buffer. 1:10 dilutions were subsequently made up to 1:10^8^ dilutions. For each dilution, we added 1 mL of the sample to each of the 3M™ Petrifilm™ (AC or Y&M) plates and distributed them evenly using the plate spreaders provided. AC plates were incubated at 35 °C for 48 h. Y&M plates were incubated at 25 °C for 5 days. Unless otherwise stated, the estimation of microbial contamination was done by the naked eye. Sample dilutions and microbial plates were prepared in the safety cabinet to minimize cross-contamination.

### 4.6 NFC-tag design and fabrication

The disposable NFC-tag comprises a flexible Printed Circuit Board (fPCB) and Android app for data visualization and communication with a cloud server. The fPCB (5.7 cm x 1.8 cm) was designed in KiCAD and manufactured by JiaLiChuang Co. Ltd to enable data acquisition and wireless transmission to a mobile device through NFC communication. An SIC4341 chip with potentiostat sensor interface and NFC capacity was provided by Silicon Craft Technology PLC. The NFC capacity was used to harvest energy from and communicate with the user’s smartphone. The passive components (2 x 0.1µF capacitors) were obtained from Digi-Key Electronics. 3M^TM^ Electrically Conductive Adhesive Transfer Tape 9703 was used to mount the PEGS on the fPCB. The fPCB with the sensor was attached to the 1L food containers or PE zip bags (24 cm x 25 cm) with polyurethane-based tattoo film (8 cm x 5 cm). A sensor window (1.7 cm x 1.3 cm) was designed in the fPCB to enable gas and water vapor to reach the surface of the sensors through the tattoo film.

## Acknowledgements

The authors would like to thank the Department of Bioengineering at Imperial College London. F.G. and L.G.-M. thank the US Army (grant number W911QY-20-R-0022) and the European Union’s Horizon 2020 research and innovation program under the Marie Sklodowska-Curie grant agreement No 101025390. F.G. and J.H. thank the EPSRC (EP/L016702/1). F.G. and J.H. would like to acknowledge Imperial College Centre for Processable Electronics and the Centre for Doctoral Training in Plastic Electronics. H.S.L thanks London Interdisciplinary Social Science Doctoral Training Partnership.

## Conflict of Interest

F.G. is a shareholder of Blakbear Ltd. G.B. is an operating member of BlakBear Ltd.

