## Supplementary Information for "Enabling Packaging Integration of Paper-based Electrical Gas Sensors for the Monitoring of Spoilage in Fresh Spinach"

**Table S1.** Material specifications for the membranes used to encapsulate PEGS.

| Commercial name | Short name | Material | Thickness (mm) | Biocompatibility |
|---|---|---|---|---|
| POREX® Porous PTFE Medical Materials MD25 | MD25 | PTFE | 0.19 | Use for medical applications [1] |
| Nitto TEMISH® porous PTFE S-NTF8031J | TEMISH | PTFE | 0.13 | Applications: medical face masks, respirators, air purifiers [2] |
| 3M™ Medical Film 9832F, Polyurethane | PU | Polyurethane | 0.02 | Suitable for wound care dressings and wearable devices [3] |
| Biaxially oriented polyester (OPET) film (OCLF) | mPET | PET | 0.0127 | Food packaging industry [4] |
| Cellulose-based compostable sealing film | Cellulose | Cellulose | -- | Food packaging industry [4] |
| Polyurethane-based tattoo film | Tattoo | Polyurethane | -- | Tattoo Wrap Waterproof Wound Antibacterial Transparent Bandage [5] |

[1]MD25 – POREX Virtek® PTFE Hydrophobic Medical Venting Porous Membrane Sheets, https://www.porex.com/product/porex-virtek-ptfe-hydrophobic-medical-venting-porous-membrane-sheets-md25/; [2] https://www.nitto.com/eu/en/products/temish_search/about/; [3]https://www.3m.co.uk/3M/en_GB/p/d/v000266868/; [4]Bullseye Food Packaging https://www.bfpuk.com/; 5 Patrick F. Mcclernon, Russell Blette, Donna Dearinger, *Transparent breathable polyurethane film for tattoo aftercare and method*, WO 2010/042511 A1.

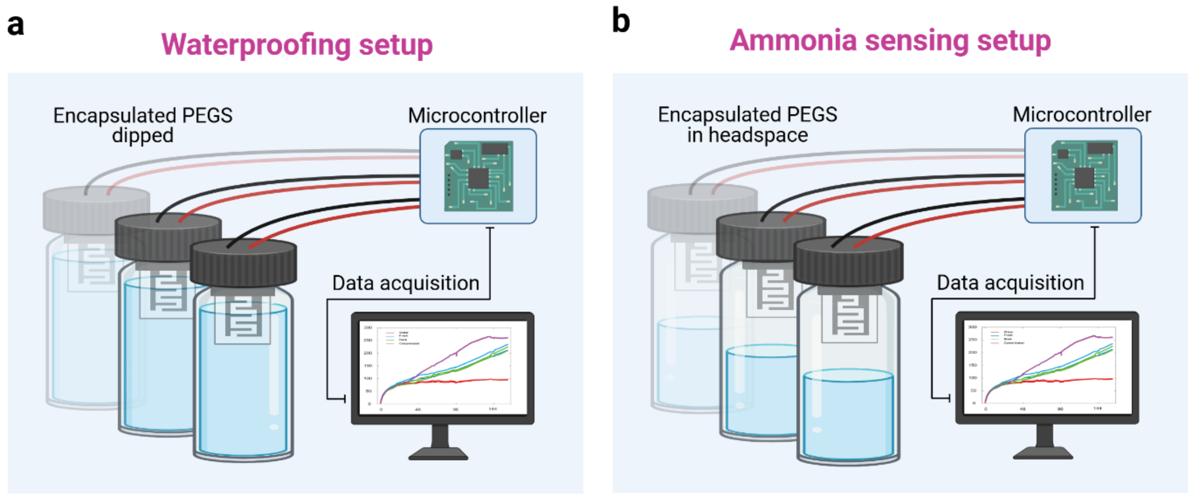


**Figure S1**. **a** Scheme of experimental setup used to evaluate the waterproofing properties of the encapsulation membranes (PEGS dipped into DI water). **b** Scheme of experimental setup used to measure changes in conductance over time of encapsulated PEGS placed in the headspace of a vial containing 1 mM $NH_4OH$ solution and comparison to non-encapsulated PEGS response.

**Table S2.** Breakdown of the cost for the fabrication of the NFC-enabled system integrated with encapsulated PEGS (Tattoo film) for the monitoring of spoilage in spinach

| Component | Price per unit (USD) |
|---|---|
| PEGS | 0.02 |
| 3M 9703 Conductive Tape | 0.06 |
| SIC4341 Chip | 0.01 |
| Passive components (capacitors) | 0.01 |
| Flexible PCB | 0.16 |
| Tattoo film | 0.08 |
| **TOTAL (USD)** | **0.35** |
